# MetaPathPredict: A machine learning-based tool for predicting metabolic modules in incomplete bacterial genomes

**DOI:** 10.1101/2022.12.21.521254

**Authors:** D. Geller-McGrath, Kishori M. Konwar, V.P. Edgcomb, M. Pachiadaki, J. W. Roddy, T. J. Wheeler, J. E. McDermott

## Abstract

The reconstruction of complete microbial metabolic pathways using ‘omics data from environmental samples remains challenging. Computational pipelines for pathway reconstruction that utilize machine learning methods to predict the presence or absence of KEGG modules in incomplete genomes are lacking. Here, we present MetaPathPredict, a software tool that incorporates machine learning models to predict the presence of complete KEGG modules within bacterial genomic datasets. Using gene annotation data and information from KEGG module databases, MetaPathPredict employs neural network and XGBoost stacked ensemble models to reconstruct and predict the presence of KEGG modules in a genome. MetaPathPredict can be used as a command line tool or as an R package, and both options are designed to be run locally or on a compute cluster. In our benchmarks, MetaPathPredict makes robust predictions of KEGG module presence within highly incomplete genomes.

## Introduction

Microbial communities play a key role in all major biogeochemical cycles on Earth. Accurate and complete identification of microbial metabolic pathways within genomic data is crucial to understanding the function of microorganisms. This identification of pathways within genomic data, and assessment of their expression, provides important insight into their influence on the chemistry of their environment and their mediation of interactions with other organisms.

In recent decades, the scientific community has significantly advanced its capability to gather and sequence genomes from microorganisms. Key steps in the process of working with isolated genomes, single-amplified genomes (SAGs), or metagenome assembled genomes (MAGs), are identifying genes coding for enzymes that catalyze metabolic reactions and inferring the metabolic potential of the associated organism from these data. These analyses involve comparing protein-coding sequences with homologous sequences from reference metabolic pathway databases including KEGG (Kanehisa et al. 2000) and MetaCyc (Caspi et al. 2018). Environmental genomes that are recovered from high-throughput sequencing samples vary in their degree of completeness due to numerous factors including limited coverage of low-abundance microbes, composition-based coverage biases, insertion-deletion errors, and substitution errors (Browne et al. 2020). Enzymes encoded in genomes are also missed due to limitations in protein annotation methods, that is, undiscovered protein families may be undetected by traditional homology-based methods. This can limit the ability to determine the extent to which these organisms (or communities) can catalyze metabolic reactions and form pathways.

Sequencing biases, novel protein families, and incomplete gene and protein annotation databases lead to missing, ambiguous, or inaccurate gene annotations that create incomplete metabolic networks in recovered environmental genomes. Existing algorithms for identifying incomplete reactions (“filling gaps”), in metabolic networks largely fall into two categories of approaches: those based on network topology (such as the method utilized by Gapseq (Zimmerman et al. 2021)), and those that utilize pre-defined pathway cutoffs (such as those used by METABOLIC (Zhou et al. 2022)). Network topology and pathway gene presence/absence cutoffs, however, can lead to underestimation of pathways that are present, particularly in highly incomplete genomes. Parsimony-based algorithms such as MinPath detect gaps in a metabolic network and identify the minimum number of modifications to the network that can be made to activate those reactions (Ye et al. 2009); its conservative approach, however, can lead to underestimation of the metabolic pathways present in a sample. Other modern tools, such as DRAM (Shaffer et al. 2020), provide annotations for metagenomic sequences but do not closely tie these to metabolic pathways. Flux-balance analysis (e.g. Escher-FBA [Rowe et al. 2018]) utilizes genome-scale metabolic models of organisms and requires experimental growth data for model parameterization; it is not easily applied to incomplete genome data, and the additional required experimental measurements may prohibit application in many use cases.

An emerging set of methods utilize machine learning models to a related problem of classifying microbial organisms’ niches based on their genomic features. One such example is a tool called Traitar, which utilizes Support Vector Machines (SVMs) to predict lifestyle and pathogenic traits in prokaryotes based on gene family abundance profiles (Weiman et al. 2016). Other recent approaches have used machine learning approaches to train models using eukaryote MAG and transcriptome data to classify trophic mode (autotroph, mixotroph, or heterotroph) based on gene family abundance profiles (Lambert et al. 2021, Alexander et al. 2021). To our knowledge, there are no existing tools that predict the presence/absence of KEGG metabolic modules (with gap-filling options) via machine learning models trained on gene features of high-quality genomes.

Here, we present “MetaPathPredict”, an open-source tool for metabolic pathway prediction based on a XGBoost (Chen and Guestrin, 2016) and neural network stacked ensemble classification framework. MetaPathPredict addresses critical deficiencies in existing metabolic pathway reconstruction tools that limit the utility and predictive power of ‘omics data: it connects manually curated, current knowledge of metabolic pathways from the KEGG database with machine learning methods to reconstruct and predict the presence or absence of KEGG metabolic modules within genomic datasets including bacterial isolate genomes, MAGs, and SAGs.

The models underlying MetaPathPredict contain metabolic reaction and pathway information from taxonomically diverse bacterial isolate genomes and MAGs from the NCBI RefSeq (O’Leary et al. 2016) and Genome Taxonomy Database (GTDB, Parks et al. 2021) databases. The set of metabolic modules from the KEGG database is the basis of the tool’s metabolic module reconstruction and prediction. The KEGG database contains metabolic pathway information for thousands of prokaryotic species and strains. KEGG modules are functional units of metabolic pathways composed of sets of ordered reaction steps. Examples include carbon fixation pathways, nitrification, biosynthesis of vitamins, and transporters or two-component systems (see Supplementary Table 1 for a description of the distribution of modules covered by MetaPathPredict). MetaPathPredict is designed to run on the command line locally or on a computing cluster and is available as an R package from GitHub (https://github.com/d-mcgrath/MetaPathPredict).

A detailed overview of the MetaPathPredict pipeline is provided in Figure 1. The tool accepts as input gene annotations (of one or more genomes) with associated KEGG ortholog (KO) gene identifiers and first reconstructs both complete and incomplete KEGG metabolic modules, then predicts the presence or absence of all incomplete modules (or modules specified by the user). Annotations can be used as input from tools such as KofamScan (Aramaki et al. 2020), DRAM, blastKOALA (Kanehisa et al. 2016), ghostKOALA (Kanehisa et al. 2016), or a custom list of KO identifier gene annotations. MetaPathPredict utilizes a stacked ensemble binary classification approach to predict the presence or absence of complete KEGG modules in incomplete genomes based on the KO gene identifiers of input genomes. MetaPathPredict classification models produced accurate results on held-out test genome annotation datasets even when the data were highly incomplete. A set of 293 neural network/XGBoost stacked ensemble models, one for each of 293 KEGG modules, made predictions with a high degree of precision on all test datasets and with high recall on genomes with an estimated completeness as low as 35%. False positive predictions were rare in all tests, while false negatives increased when predictions were made with highly incomplete gene annotation information, as would be expected. An additional set of 183 neural network models (not stacked ensembles) trained to classify the presence of KEGG modules rarely present in the training data made predictions with robust precision and recall on genomes with a completeness estimate of at least 60%, exhibiting very similar precision and recall characteristics to the stacked ensembles. We believe that MetaPathPredict will be a valuable tool to enable studies of metabolic potential in environmental microbiome studies as well as synthetic biology effort.

**Figure 1.**
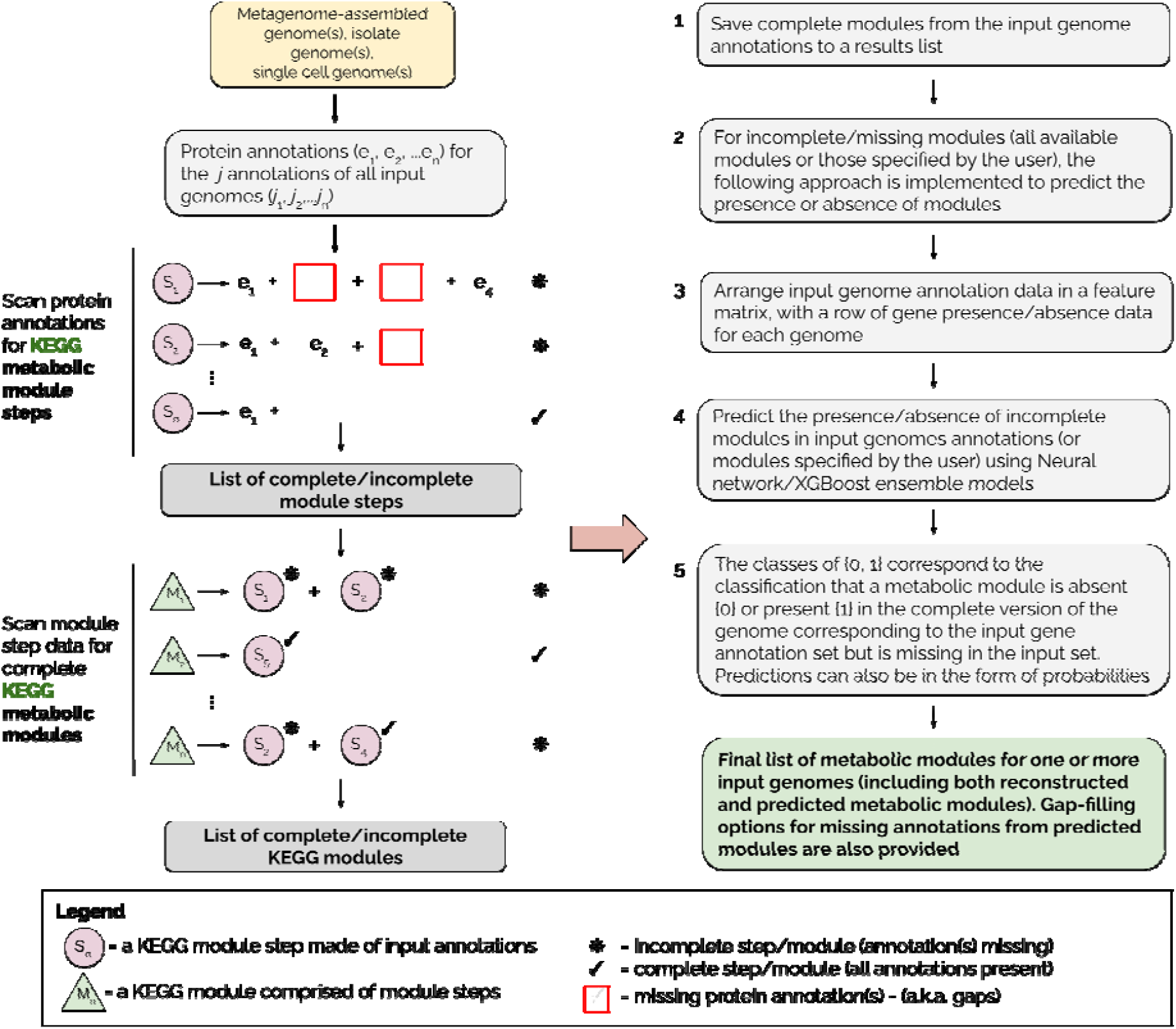
Overview of the MetaPathPredict pipeline. Input genome annotations are read into MetaPathPredict as a data object. The data are scanned for present KEGG modules and are formatted into a feature matrix. The feature matrix is then used to make predictions for all incomplete modules (or modules specified by the user). A summary and detailed reconstruction and prediction objects, along with gap-filling options are returned in a list as the final output.

## Results and Discussion

MetaPathPredict’s performance metrics on held-out test datasets suggest its models predict with high fidelity when at least 40% of gene annotations are recovered from a reconstructed genome (Figure 2). The efficacy of MetaPathPredict models was assessed using incomplete gene annotation data simulated from whole genomes, as well as from genomes reconstructed from reads that had been randomly downsampled. We further benchmarked MetaPathPredict against custom presence/absence classification rules, and existing gap-filling tools METABOLIC and Gapseq.

**Figure 2.**
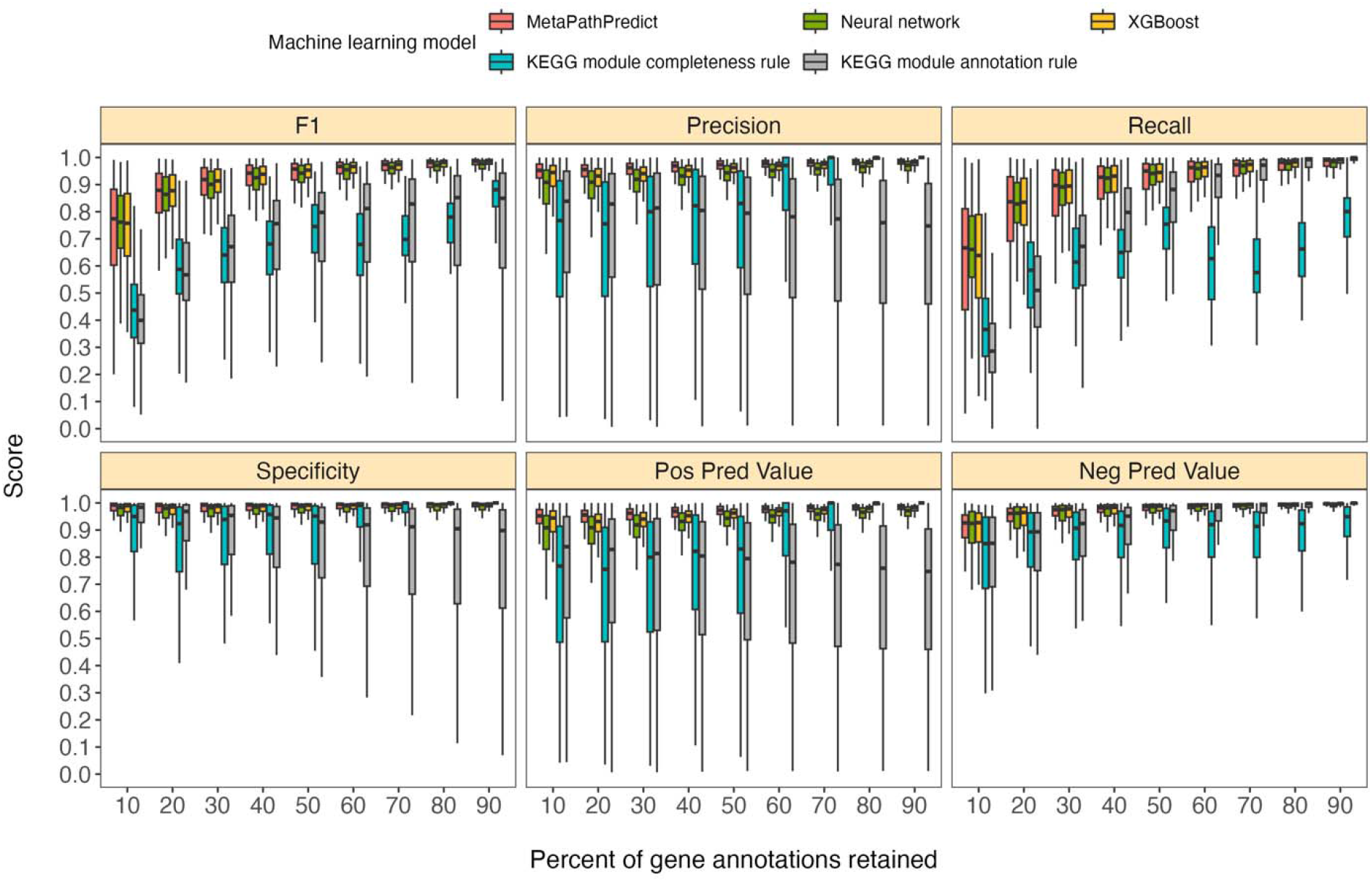
Comparison of performance metrics of MetaPathPredict’s 293 stacked ensemble binary classification models to next-best performing XGBoost and neural network machine learning models as well as two naïve classification rules. Downsampled gene annotations of high-quality genomes used in held-out test sets are from NCBI RefSeq and GTDB databases. Each boxplot displays the distribution of model performance metrics for predictions on randomly sampled versions of the gene annotation test sets in downsampling increments of 10% (90% down to 10%, from right to left). The binary classifier performances are based on the classification of the presence or absence of KEGG modules in the complete versions of the gene annotations that were downsampled for model testing.

### Benchmarking MetaPathPredict and alternative strategies on downsampled NCBI RefSeq and GTDB data

We compared the performance of MetaPathPredict’s stacked ensemble models to two classes of competitor classifiers: naive rule-based classifiers and various other machine learning model architectures. The evaluation was performed on test datasets comprised of isolate and metagenome-assembled genomes from the GTDB and NCBI databases (see Methods). When evaluating with the same sets of randomly downsampled gene annotations, we found that each competing method performed worse than MetaPathPredict (Figure 2). We assessed two naïve classification methods. First, we devised a classification rule based on the completeness of a KEGG module relative to the number of genes retained after downsampling: if, in a downsampled genome, the number of genes involved in a KEGG module are present in at least the same proportion of all genes retained, the KEGG module is classified as ‘present,’ otherwise it is labeled ‘absent’. For example, if 50% of gene annotations were removed from a genome during downsampling, then any KEGG module for which 50% of its associated genes are retained would be reported as ‘present’. The results of this naive approach (Figure 2) show that the relative completeness of a KEGG module alone is not a robust classification strategy. The second classification rule that we tested was: for all gene annotation sets in the dataset, if any genes were present in an annotation set that were unique to a KEGG module (relative to other modules) then the module was classified as ‘present’, otherwise it was labelled ‘absent’. The results of this naive approach (Figure 2) suggest that the presence of rare protein annotations or genes unique to a certain KEGG module is not always a strong indicator of the presence of a module in a genome. Ultimately, the performance of these naive classifiers indicate that MetaPathPredict’s models have the advantage of incorporating information from genes outside of KEGG modules. We additionally compared the performance of various machine learning model architectures. Of these, the XGBoost (no ensemble) and neural network (no ensemble) architectures were the next-best performing models (Figure 2).

MetaPathPredict’s ensemble strategy produced the best observed performance. Mean F1 score of the models was at least 0.90 when predicting on test datasets in which 30-90% of gene annotations had been retained. MetaPathPredict rarely made false positive predictions based on data from highly incomplete gene annotation sets; the average precision of the models was consistently above 0.94 for all held-out test sets. MetaPathPredict also did not misclassify most negative class observations; the 293 models had an overall average specificity above 0.9 on all test datasets. The quality of MetaPathPredict’s negative class predictions was robust on test datasets containing at least 40% of the complete gene annotation data. The mean recall decreased incrementally with decreased gene annotation data and was below 0.86 on only on test sets containing 30% or less of the complete gene annotation data. The drop in recall and negative predictive value on highly downsampled test data was indicative of an increase in false negative predictions when MetaPathPredict models were assessing KEGG module presence/absence based on highly incomplete information.

### Benchmarking MetaPathPredict against GEM repository MAGs

MetaPathPredict was further tested on gene annotations from a set of 100 high-quality metagenome-assembled genomes from the GEM genome repository (Figure 3.). The MAGs selected from the repository had an estimated completeness of 100 and estimated contamination of 0, and MIMAG quality score of “High Quality”. The genomes belonged to 15 taxonomic classes and were recovered from 11 different environments, primarily from human-associated and built environment metagenomes (see Supplementary Fig. 4 for GEM genome taxonomic distributions and environmental sources). We created a list of 9 GEM datasets by randomly downsampling the data to retain 10% to 90% of gene annotations (in 10% increments) as in the previous section. MetaPathPredict classified the presence/absence of KEGG modules in each MAG. Overall results were comparable to MetaPathPredict’s performance on the benchmark based on GTDB and NCBI databases. The models excelled at predicting the presence or absence of KEGG modules in genomes when at least 50% of gene annotations were randomly retained. Predictions were less reliable when 40% or less gene annotation data was retained.

**Figure 3.**
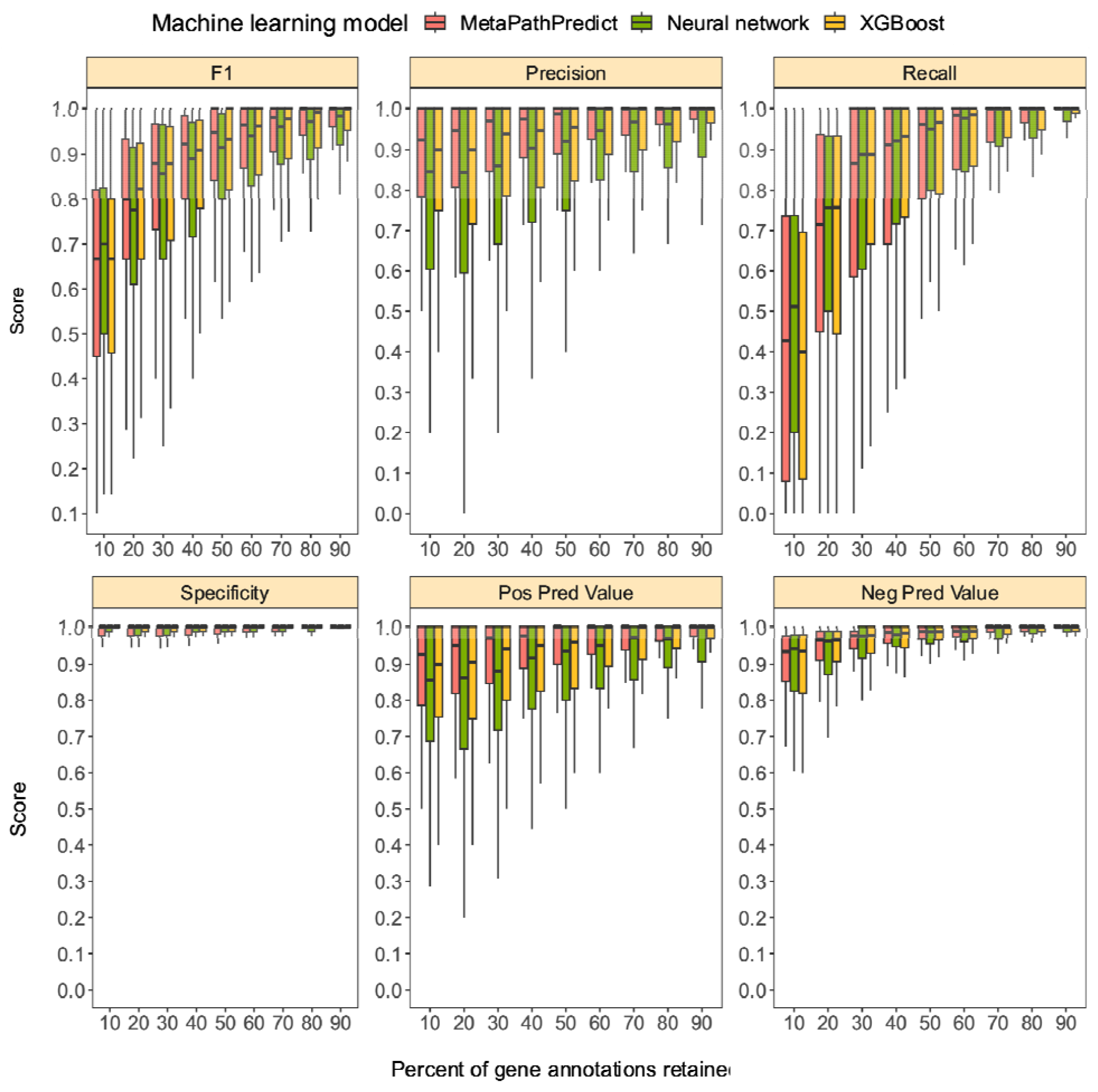
Boxplots of performance metrics of 293 MetaPathPredict models on high-quality bacterial GEM MAGs (*n* = 100). Model performance metrics are for predictions on downsampled versions of GEM genome gene annotations in decreasing increments of 10% (retaining 10-90% of the annotations in each test set). MetaPathPredict’s stacked ensemble models were benchmarked against XGBoost and neural network model architectures.

### Benchmarking MetaPathPredict against existing tools on a deataset with downsampled reads

In addition to model assessments made through downsampling protein annotations, we evaluated a second set of held-out test set genomes (*n* = 50) that we downsampled at the sequence read level. We downsampled genome sequence reads to assemble and annotate fragmented versions of these test set genomes from NCBI and GTDB. MetaPathPredict’s performance on this test set closely resembled protein annotation random sampling results (Figure 4a, Figure 4b).

**Figure 4.**
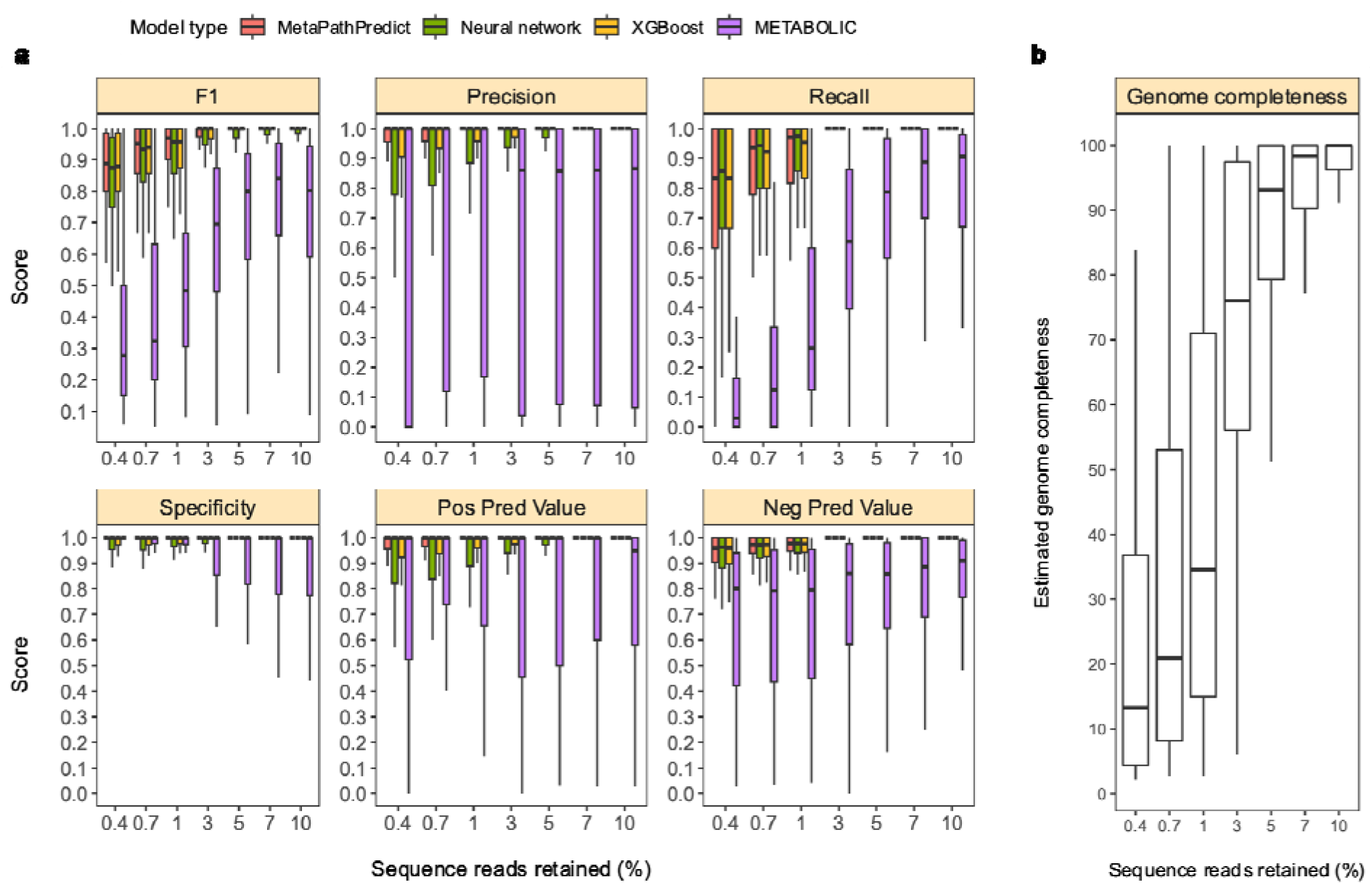
Performance metrics boxplots of 293 stacked ensemble classification models. Downsampled sequence reads of high-quality genomes used as a second held-out test set are from NCBI RefSeq and GTDB databases. **Panel a:** Boxplots display the distribution of model performance metrics for predictions of KEGG module presence/absence on simulated incomplete genomes downsampled at the sequence read level by MetaPathPredict models and METABOLIC. Downsampling increments were chosen based on average estimated completeness of the test set genomes at each increment to reflect a range of estimated completeness thresholds. **Panel b:** Average estimated genome completeness distributions of test set genomes that were downsampled at the sequence read level and then assembled and annotated.

MetaPathPredict had an average F1 score for all 293 models of 0.92 on the second held-out test sets that had an average estimated genome completeness of 35%. The similarity of these results to the gene annotation downsampling approach validates the latter approach that was used more broadly to assess MetaPathPredict. We compared MetaPathPredict’s performance to module presence/absence predictions made on the same data with METABOLIC, which is a command line tool that performs gene annotations and estimates the completeness of KEGG Modules in individual genomes and prokaryotic microbial communities (Figure 4a). We found that MetaPathPredict’s models made much more sensitive predictions on test set genomes. We also benchmarked MetaPathPredict against Gapseq based on both tools’ predictions of the presence or absence of a particular KEGG pathway in test set genomes (Figure 5a). MetaPathPredict outperformed predictions made by Gapseq, particularly on genomes that had had a high proportion of reads removed prior to assembly.

**Figure 5.**
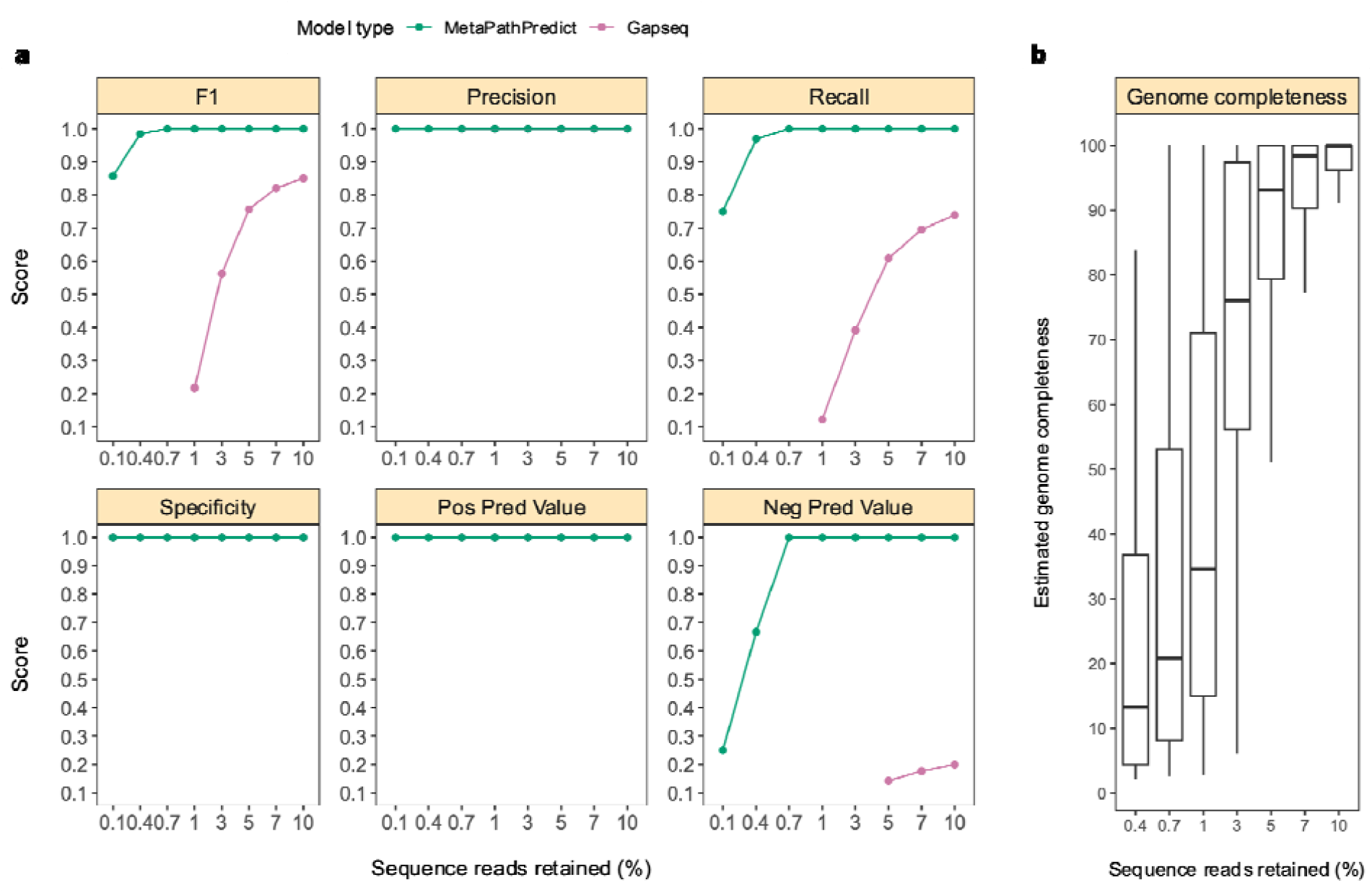
Performance metrics boxplots of MetaPathPredict and Gapseq predictions for KEGG pathway map00290 (Valine, leucine, and isoleucine biosynthesis) which contains KEGG modules M00019, M00432, M00535, and M00570. For MetaPathPredict predictions, the whole KEGG pathway was considered present if the aforementioned KEGG modules were all present. Downsampled sequence reads of high-quality genomes used as a second held-out test set are from NCBI RefSeq and GTDB databases. **Panel a:** Line segments display model performance metrics for MetaPathPredict and Gapseq predictions of KEGG pathway map00290 presence/absence on simulated incomplete genomes downsampled at the sequence read level. Downsampling increments were chosen based on average estimated completeness of the test set genomes at each increment to reflect a range of estimated completeness thresholds. **Panel b:** Average estimated genome completeness distributions of test set genomes that were downsampled at the sequence read level and then assembled and annotated.

We fitted and tested an additional 183 neural network models for rare KEGG modules present in less than 5% of the training data (Supplementary Figs. 2b, 5-6). The neural network models were not incorporated into ensembles with XGBoost models due to the severe class imbalance for these KEGG modules. These models performed optimally on genomes that were estimated to be at least 60% complete.

## Conclusion

MetaPathPredict is an open-source tool that can be used to characterize the functional potential of one or more sample genomes by detecting complete KEGG modules and predicting the presence or absence of those that are incomplete or missing. The tool accepts sets of gene annotations of individual genomes in KO gene identifier format as input. This type of annotation format can be acquired by annotating a genome of interest using KEGG-based annotation tools such as KofamScan (Aramaki et al., 2020), blastKOALA (Kanehisa et al. 2016), or ghostKOALA (Kanehisa et al. 2016). MetaPathPredict also provides gene gap-filling options by suggesting putative KO gene annotations that could fill in missing gaps in predicted modules.

MetaPathPredict further validates the use of gene copy frequency within a genome as a feature for bacterial metabolic function predictions. The performance metrics of MetaPathPredict on NCBI/GTDB and GEM test datasets validated the use of stacked ensemble models to predict the presence/absence of KEGG metabolic modules with high fidelity on sparse to near-complete bacterial genomes. MetaPathPredict’s binary classification models consistently made predictions with high precision, recall and specificity on simulated and real genomes using gene annotation and sequence read downsampling methods. The predictive power of the stacked ensemble models was most limited when predicting on 10%-30% of protein annotations, and when the mean estimated completeness of reconstructed genomes from downsampled reads was below 35%. We suggest that optimal performance with MetaPathPredict can be achieved when at least 40% of genes have been recovered in an input bacterial gene annotation dataset.

MetaPathPredict will facilitate more complete and accurate reconstruction of the metabolic potential encoded within bacterial genomes from a diverse array of environments and will enhance the ability to infer what metabolisms they are capable of, and/or how they may respond to perturbations. MetaPathPredict connects the field of machine learning with the growing community of environmental microbiologists using genomic sequencing techniques and will help transform and improve the way they work with environmental genomic datasets.

## Materials and Methods

### Filtering genome database metadata, downloading high-quality genomes, and gene annotation

The NCBI RefSeq (Release 205) database metadata file was downloaded and filtered to retain only the information for all bacterial genomes classified as “Complete genome”, which are defined on the NCBI assembly help webpage “all chromosomes are gapless and have no runs of 10 or more ambiguous bases (Ns), there are no unplaced or unlocalized scaffolds, and all the expected chromosomes are present (i.e. the assembly is not noted as having partial genome representation). Plasmids and organelles may or may not be included in the assembly but if present, then the sequences are gapless.” This resulted in 17,491 complete NCBI genomes.

The GTDB bacterial metadata file for release 95 was downloaded and filtered to keep the information for all genomes with an estimated completeness greater than 99, an estimated contamination of 0, and a MIMAG (Bowers et al., 2017) quality score of “High Quality”. A total of 30,760 non-redundant bacterial genomes from the two database metadata files were downloaded using the ncbi-genome-download command line tool (Blin, K.). The RefSeq genomes (17,491 total) were downloaded from the RefSeq FTP server, and the GTDB genomes (13,269 total) were downloaded from the GenBank FTP server (Supplementary Figure 1). Genes were called using Prodigal (Hyatt et al., 2010), and the KofamScan command line tool (Aramaki et al. 2020) was used to generate gene annotations (in KO gene identifier format) for all of the genomes using the KOfam set of Hidden Markov Models (HMMs) available for download from the KEGG database (Kanehisa et al. 2002).

### Formatting gene annotation data, fitting KEGG module classification models

The gene copy number data of the downloaded genomes was formatted in a matrix containing KO gene identifier counts in columns and genomes in rows. The label of each model was the presence/absence of a KEGG module, as was determined using the KEGG modules downloaded from the KEGG database. For the module to be categorized as present, every step of the module had to be present in a genome, otherwise it was designated as absent. 293 KEGG modules were individually modelled using the gene annotation data of the genomes consolidated from the NCBI and GTDB databases. All 293 models were constructed for KEGG modules for which at least 5% of the training genomes contained the module genes. The constructed models classify the presence or absence of complete KEGG modules based on the gene annotations of a genome. Training and test sets contained both complete and incomplete gene annotations of bacterial genomes from a diverse array of phyla (Supplementary Fig. 1). The incomplete annotations used in training and testing of MetaPathPredict’s models were constructed from complete genome annotations that were randomly downsampled to retain 10-90% of the total gene content. The training datasets had a size of 24,517 observations, for which one observation was a vector in which each element was a count of the frequency that an individual KO identifier was annotated in a genome. The test datasets each contained 6,129 observations. The percent of observations with a positive class (a complete KEGG module ‘present’ in the gene annotations) in the training and test datasets varied, with a mean of 28.24% (Supplementary Fig. 2a).

A neural network and XGBoost stacked ensemble classification approach was chosen to model the relationship between whole genome KO gene identifier annotation data and the presence of metabolic modules. Model stacking is an approach in which several independently-trained models (often comprised of multiple machine learning architectures) are applied to an input, and their output is passed to another machine learning model (a ‘meta-learner’), which produces a final classification. The stacking approach in MetaPathPredict employs an L1-regularized linear model as a meta-learner to learn how to best combine predictions from the ensemble of models. Training data was divided into a subset used to train ensemble members (24,517 observations) and the remaining entries (so called ‘out-of-fold’ predictions) held-out for meta-learner training. The same training data was used to train each of the ensemble models, and the meta-learner was trained on the out-of-fold predictions made by candidate ensemble members. L1-regularization was utilized by the meta-learner to pick the final number of ensemble members from the pool of candidates. For each model trained, the meta-learner then retained ensemble members with the best performance on out-of-fold data and assigned weights to the predictions made by each ensemble member for future predictions on unseen data. Features used in each training dataset for classification were the gene copy number for all genes that were assigned KO gene identifiers (for which one observation was a vector in which each element was a count of the times that an individual KO gene identifier was annotated in a genome). Information gain, which is a measure of the statistical dependence between two variables, was utilized for feature selection to decrease model complexity and avoid over-fitting using the recipeselectors R package (Steven Pawley, 2022). This method iteratively calculates the mutual information between the dependent variable (KEGG module presence/absence) and each independent variable (the KO gene identifiers) and measures the reduction in uncertainty for the dependent variable, given the values of the independent variable. The mutual information, *I*(*X* ; *Y*), between two random variables *X* and *Y* is: *I*(*X* ; *Y*) = *H*(*X*) - *H*(*X*|*Y*), where *H*(*X*) is the entropy for *X*, and *H*(*X*|*Y*) is the conditional entropy for *X* given *Y*. The number of best-scoring features retained in each model’s train/test datasets were chosen for each model through cross validation. XGBoost models and feed-forward neural networks with a single hidden layer were used as the machine learning architectures in MetaPathPredict’s stacked ensemble models.

The full set of available genome annotations was used to form each model’s train and test datasets. Stratified random sampling, in which the genome dataset observations were divided into strata and sampled independently, was used to generate 80% train/20% test dataset splits that each contained data observations with preserved proportions of positive (‘KEGG module present’) and negative (‘KEGG module absent’) classes that were present in the genome dataset (see boxplot of the distribution of module presence/absence classes in Supplementary Figure 2a). The Tidymodels R package (Kuhn M., Wickham H., 2020) was used to fit 293 neural network/XGBoost stacked ensemble models each corresponding to a unique KEGG module.

The 293 MetaPathPredict stacked ensemble models were fit using the Stacks R package (https://github.com/tidymodels/stacks). First, potential XGBoost and neural network parameter values were assessed using 10-fold nested cross validation to maximize the Receiver Operating Characteristic (ROC). Grid search was utilized for the XGBoost parameters *mtry, min n, tree depth, learn rate, loss reduction*, and *sample size*. After locking choices for these parameters, a separate grid of values for neural etwork parameters *hidden units, penalty, dropout* and *activation* were then assessed. The Stacks package functions were then utilized to fit Least Absolute Shrinkage and Selection Operator (LASSO) meta-learner models to out-of-fold predictions made by the XGBoost and neural network models during cross validation via grid search. This was done over a range of, *λ* values 1×10^−6^ to 1×10^−1^, and the, *λ* that maximized the ROC was chosen for each ensemble member. The *l*_1_-penalization of the LASSO model facilitated the selection of final model ensemble members from the pool of candidate members for each stacked model. For 183 KEGG modules rarely present in the training data (< 5% genomes contained the full module), neural network models were chosen based on their outperformance of XGBoost models on test datasets instead due to severe class imbalance and inability to fit stacked ensemble models to these KEGG modules.

We assessed and benchmarked MetaPathPredict’s models against two naïve classification methods. First, we hypothesized that we could predict the presence of a KEGG module if, after downsampling test sets of gene annotations, the proportion of module genes present in the dataset was greater than or equal to the percentage of annotations retained in the dataset. If the proportion of genes involved in a KEGG module were present in a dataset observation at least equivalently to the proportion of gene annotations retained after downsampling, the module was classified as ‘present’, otherwise it was classified as ‘absent’. The second naïve classification rule was: for all gene annotation sets in the dataset, if any genes were present in an annotation set that were unique to a KEGG module (relative to all other KEGG modules) then the module was classified as ‘present’, otherwise it was labelled ‘absent’. We additionally benchmarked MetaPathPredict’s stacked ensemble models against several other machine learning model types including standalone neural network and XGBoost models.

### Evaluating models on test data, including test data randomly downsampled to simulate varying degrees of genome incompleteness

Each of MetaPathPredict’s models was validated on a held-out test set with 6,129 observations and the performance metrics were extracted using the Tidymodels R package. The performance metrics used in evaluating the models were precision, recall, specificity, F1 score, positive predictive value, and negative predictive value (Table 1).

**Table 1.** Definitions of machine learning model performance metrics used to assess MetaPathPredict models. *Prevalence is defined due to its use in the definitions of negative and positive predictive value.

| Metric | Definition |
| --- | --- |
| Precision | $\text{true positive} / (\text{true positive} + \text{false positive})$ |
| Recall | $\text{true positive} / (\text{true positive} + \text{false negative})$ |
| Specificity | $\text{true negative} / (\text{true negative} + \text{false positive})$ |
| F1 score | $2 \times ((\text{precision} \times \text{recall}) / (\text{precision} + \text{recall}))$ |
| Positive predictive value | $\text{recall} \times \text{prevalence} / ((\text{recall} \times \text{prevalence}) + ((1 - \text{specificity}) \times (1 - \text{prevalence})))$ |
| Negative predictive value | $\text{specificity} \times (1 - \text{prevalence}) / ((1 - \text{recall}) \times \text{prevalence}) + (\text{specificity} \times (1 - \text{prevalence}))$ |
| Prevalence* | $(\text{true positive} + \text{false negative}) / (\text{true positive} + \text{false positive} + \text{true negative} + \text{false negative})$ |

The genome annotations from each test set were then randomly downsampled to simulate recovered gene annotations from incomplete genomes. 10% to 90% of genes from each annotation set were randomly retained (in increments of 10%) and used as input for MetaPathPredict predictions of KEGG module presence/absence. The model performance metrics in each simulation were extracted using the Tidymodels package.

### Testing models with a set of high-quality metagenome-assembled genomes from the Genomes from Earth’s Microbiomes online repository

MetaPathPredict was further validated on another test set of genome annotations extracted from the Genomes from Earth’s Microbiomes (GEM; Nayfach et al. 2021) repository of MAGs. The GEM metadata file was downloaded from the repository (Nayfach et al. 2021) and filtered to retain a random sample of 100 MAGs with a CheckM (Parks et al. 2015) estimated completeness of 100, an estimated contamination of 0, and a MIMAG quality score of “High Quality”. The method for this assessment was the same as was described above for testing MetaPathPredict model performances on the held-out test data.

### Evaluating models on test data downsampled at the read level

A second held-out set of genomes (*n* = 50) was downloaded from NCBI/GTDB databases using the SRA Toolkit (SRA Toolkit Development Team) and SRA explorer (Phil Ewels). The raw sequencing reads were filtered using fastp (Chen et al. 2018), and the quality-filtered reads were randomly downsampled using seqtk (Li, H. 2012). Downsampled reads were assembled into genomes using the SPAdes assembler (Bankevich et al., 2012), genes were called with Prodigal and then annotated using KofamScan. MetaPathPredict’s stacked ensemble models were then used to predict the presence or absence of all 293 KEGG modules in each genome and predictions were then cross-referenced with their known presence/absence based on the unmodified test dataset. In addition to simple approaches described above, the METABOLIC (Zhou et al. 2022) and Gapseq (Zimmerman et al. 2021) tools were evaluated on the same benchmark dataset. The METABOLIC-G.pl program was used with default settings. Gapseq (using default settings) and MetaPathPredict were compared by evaluating predictions for the presence or absence of the KEGG pathway map00290 (Valine, leucine, and isoleucine biosynthesis) which contains KEGG modules M00019, M00432, M00535, and M00570. For MetaPathPredict predictions, the whole KEGG pathway was considered present if the aforementioned KEGG modules were all predicted as present, otherwise it was classified as absent.

### Gapfilling for incomplete modules predicted as present

MetaPathPredict provides enzyme gap-filling options for KEGG modules predicted as present by suggesting putative KO gene annotations missing from an input genome’s gene annotations that could fill in missing gaps in predicted modules.

## Acknowledgements

D. Geller-McGrath acknowledges funding from DOE SCGR Fellowship for the 2020 Solicitation 2 in Computational Biology and Bioinformatics. We would like to thank A. Solow (WHOI) for helpful initial discussion about statistical approaches. JEM, JWR and TJW were supported by the Department of Energy (DOE) Office of Biological and Environmental Research (BER) through the “Machine-Learning Approaches for Integrating Multi-Omics Data to Expand Microbiome Annotation” project. PNNL is operated for the DOE by Battelle Memorial Institute under Contract DE-AC05-76RL01830.

## Ethics declarations

The authors declare no competing interests.

## Data Availability

Genomic data used for creation of MetaPathPredict models is available from the NCBI Bacterial RefSeq Genomes database (https://ftp.ncbi.nlm.nih.gov/genomes/refseq/bacteria/, version 209) and the Genome Taxonomy Database (https://data.gtdb.ecogenomic.org/releases/latest/, version r95). The GEM genomes used for model benchmarking are available at the GEM repository (https://portal.nersc.gov/GEM/genomes/). The sequencing reads used for model benchmarking are available at the NCBI Sequence Read Archive website (https://www.ncbi.nlm.nih.gov/sra).

## Code Availability

The scripts used for all data processing, model training, model benchmarking, and figure creation used in this study are available in the following GitHub repository: https://github.com/Microbiaki-Lab/MetaPathPredict_workflow. The MetaPathPredict R package is available from the following GitHub repository: https://github.com/d-mcgrath/MetaPathPredict.

## Supplementary Figures

**Supplementary Figure 1.**
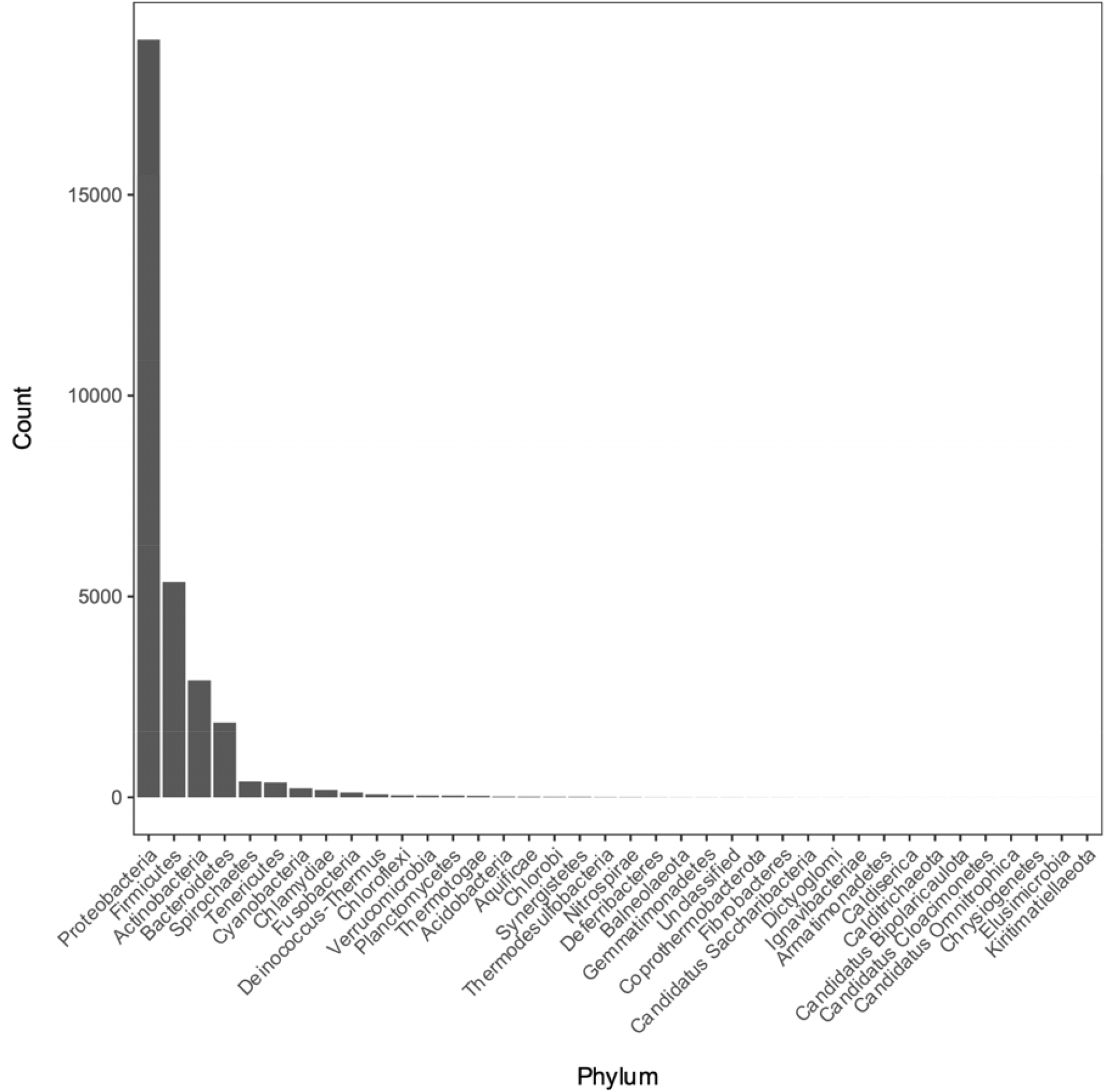
Distribution of phyla of bacterial genomes from which annotation data was used during model training and testing. See Supplementary Table 2 for the full metadata table.

**Supplementary Figure 2.**
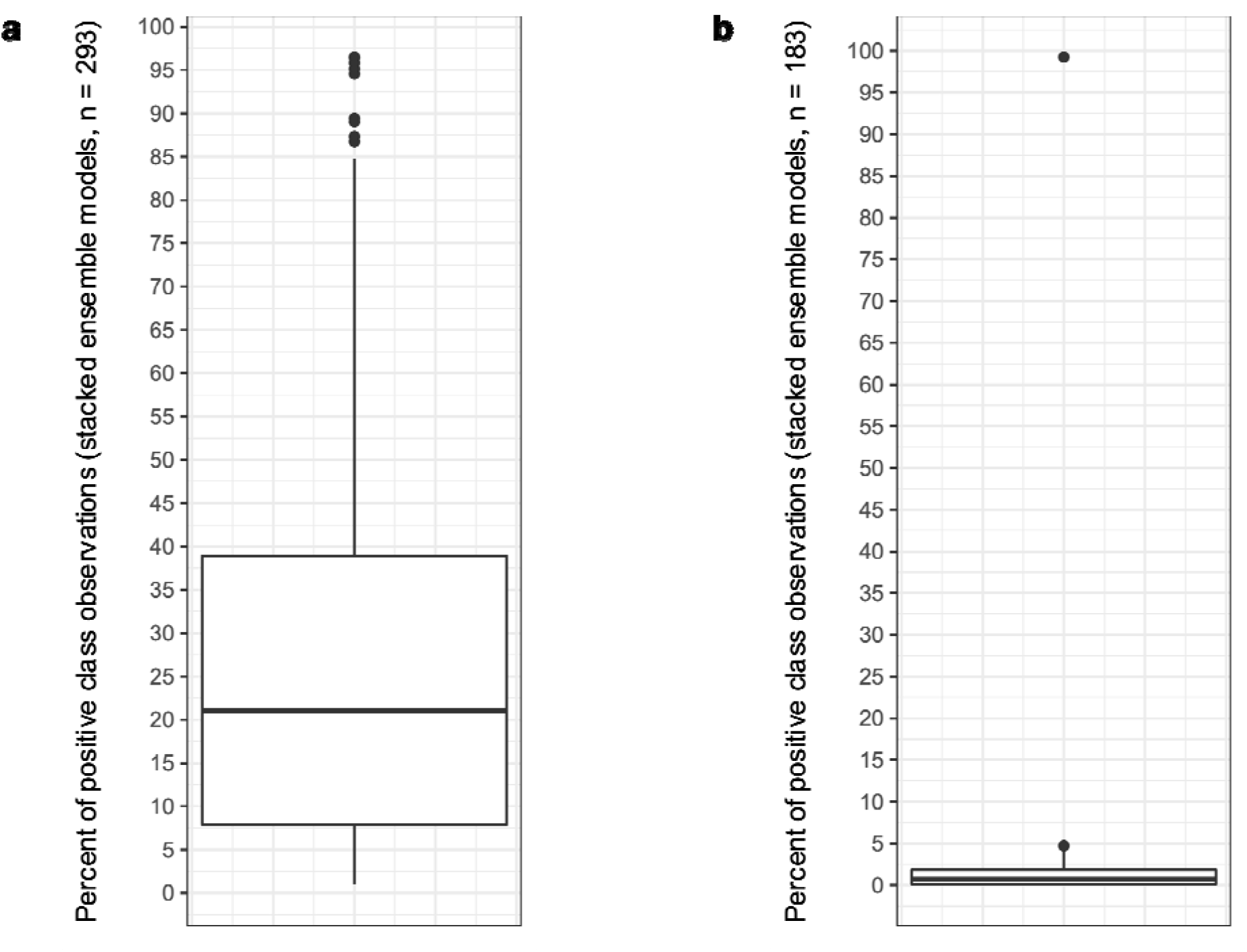
Boxplot of the ratio of positive to negative classes of observations in all MetaPathPredict stacked ensemble training and test datasets (**a**, *n* = 293) and neural network models (**b**, *n* = 183). Each train/test dataset split contains the same distribution of observation classes.

**Supplementary Figure 3.**
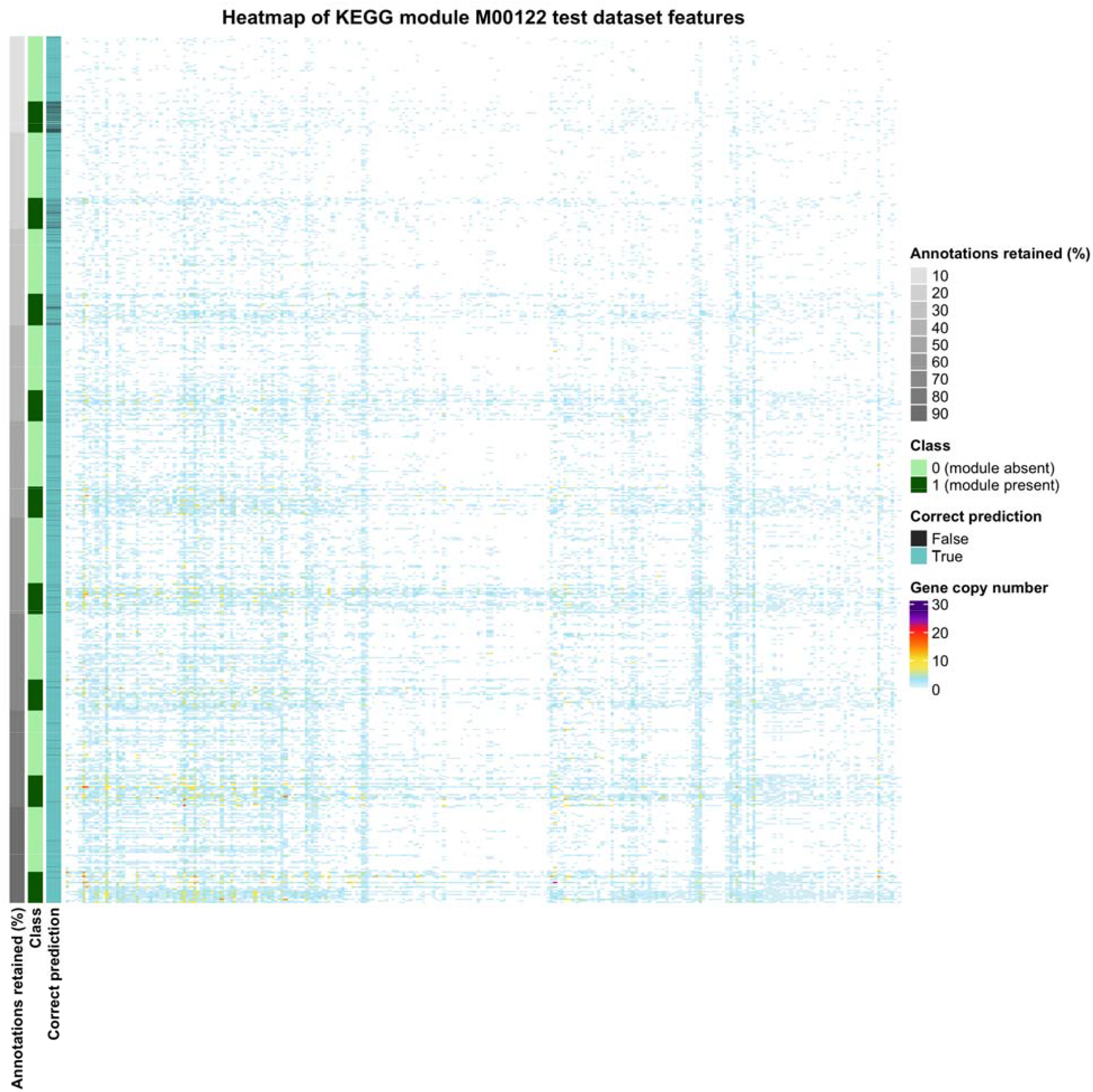
Gene copy number data for the set of features (genes) used by MetaPathPredict’s stacked ensemble model for KEGG module M00122 (cobalamin biosynthesis). This gene annotation data is from the held-out test set of downsampled protein annotation data for KEGG module M00122, and is only showing the gene copy number of genes used by the stacked ensemble model for making predictions.

**Supplementary Figure 4.**
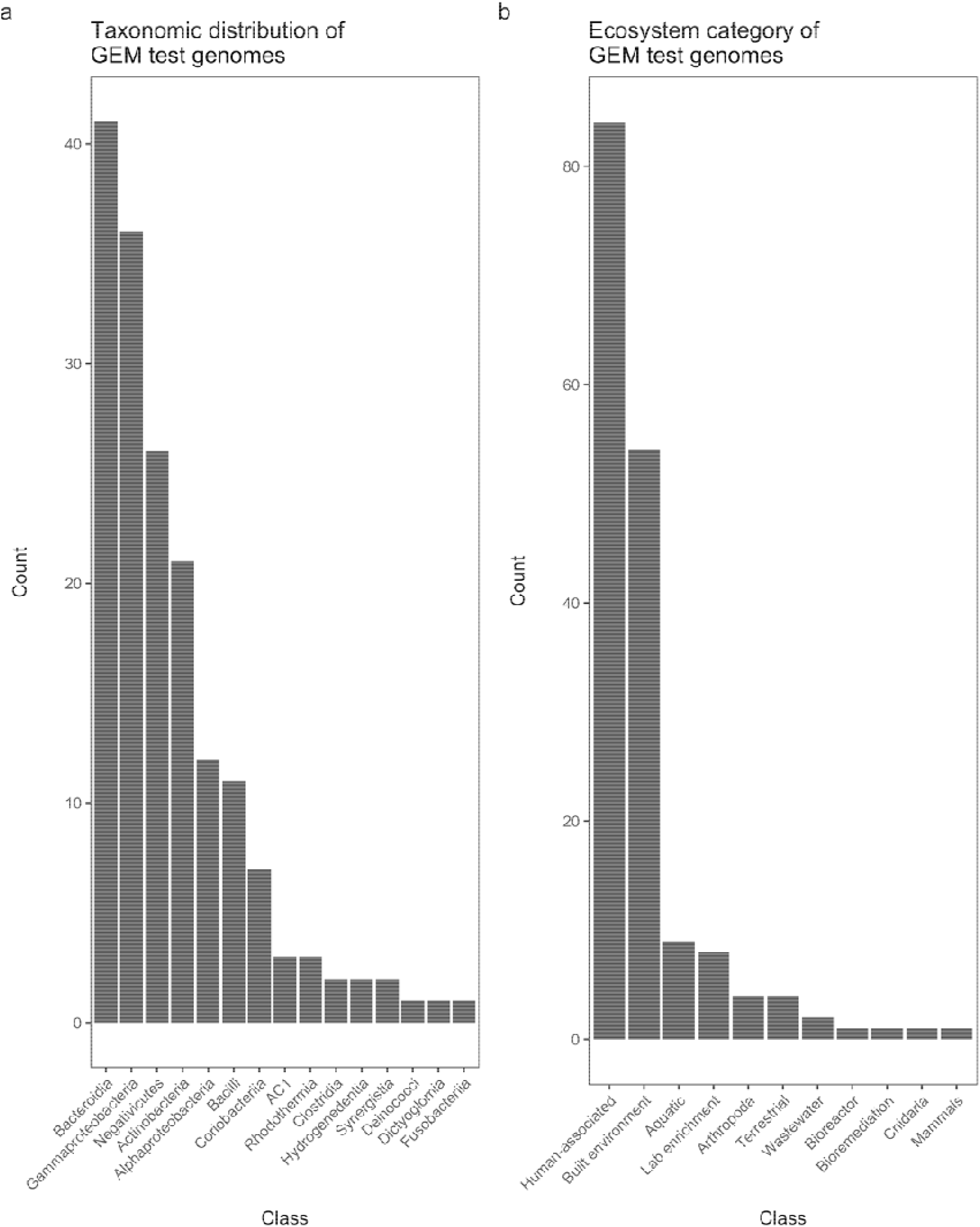
**Panel a**: Bar chart of the taxonomic distribution of genomes from the GEM repository used during model validation. **Panel b**: Bar chart of the environmental sources of metagenomes the MAGs from this test set were recovered from.

**Supplementary Figure 5.**
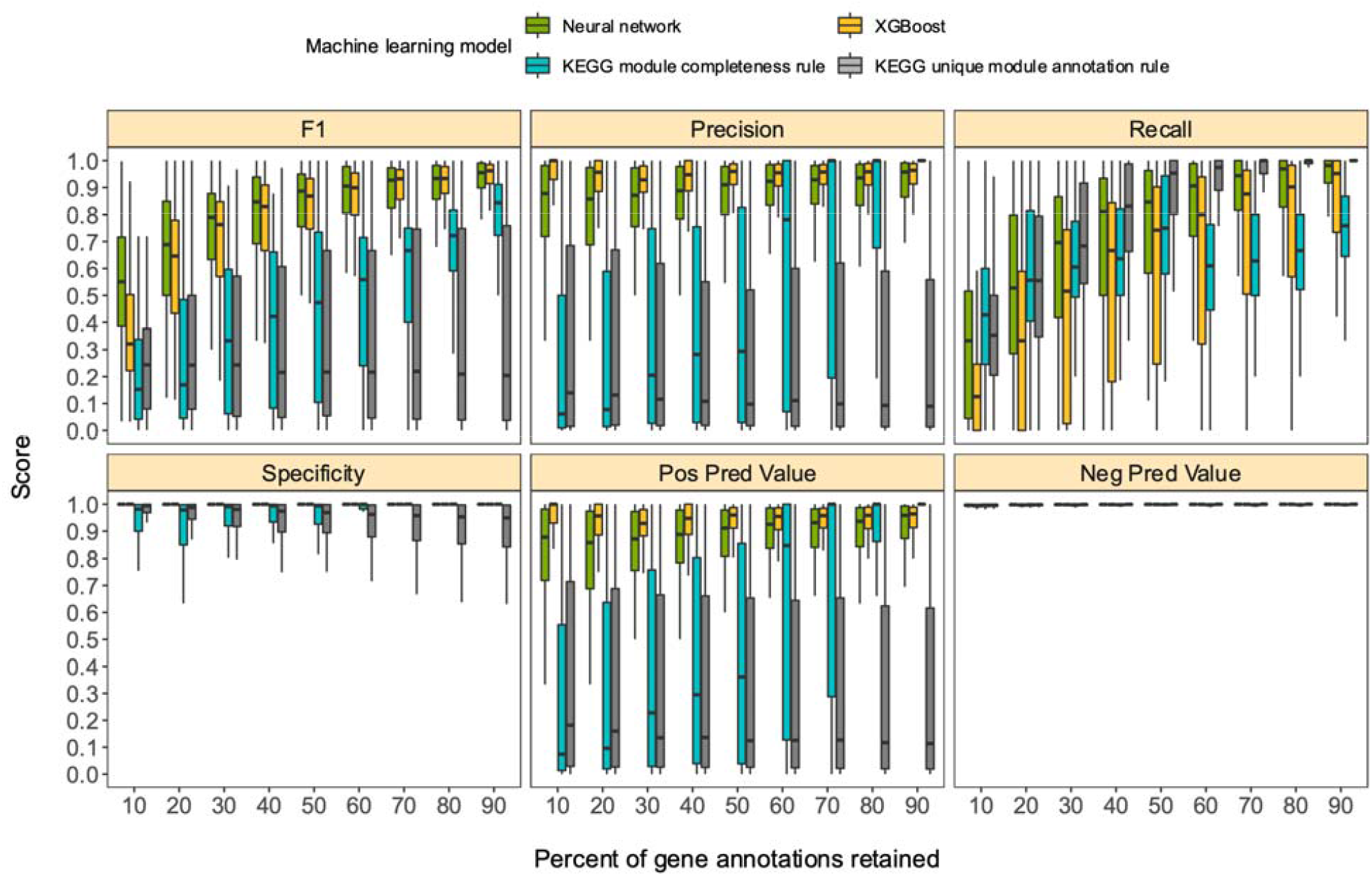
Boxplots of performance metrics of 183 neural network and XGBoost binary classification models. Downsampled gene annotations of high-quality genomes used in held-out test sets are from NCBI RefSeq and GTDB databases. Each boxplot displays the distribution of model performance metrics for predictions on randomly sampled versions of the gene annotation test sets in downsampling increments of 10% (90% down to 10%, from right to left). The binary classifier performances are based on the classification of the presence or absence of KEGG modules in the complete versions of the gene annotations that were downsampled for model testing.

**Supplementary Figure 6.**
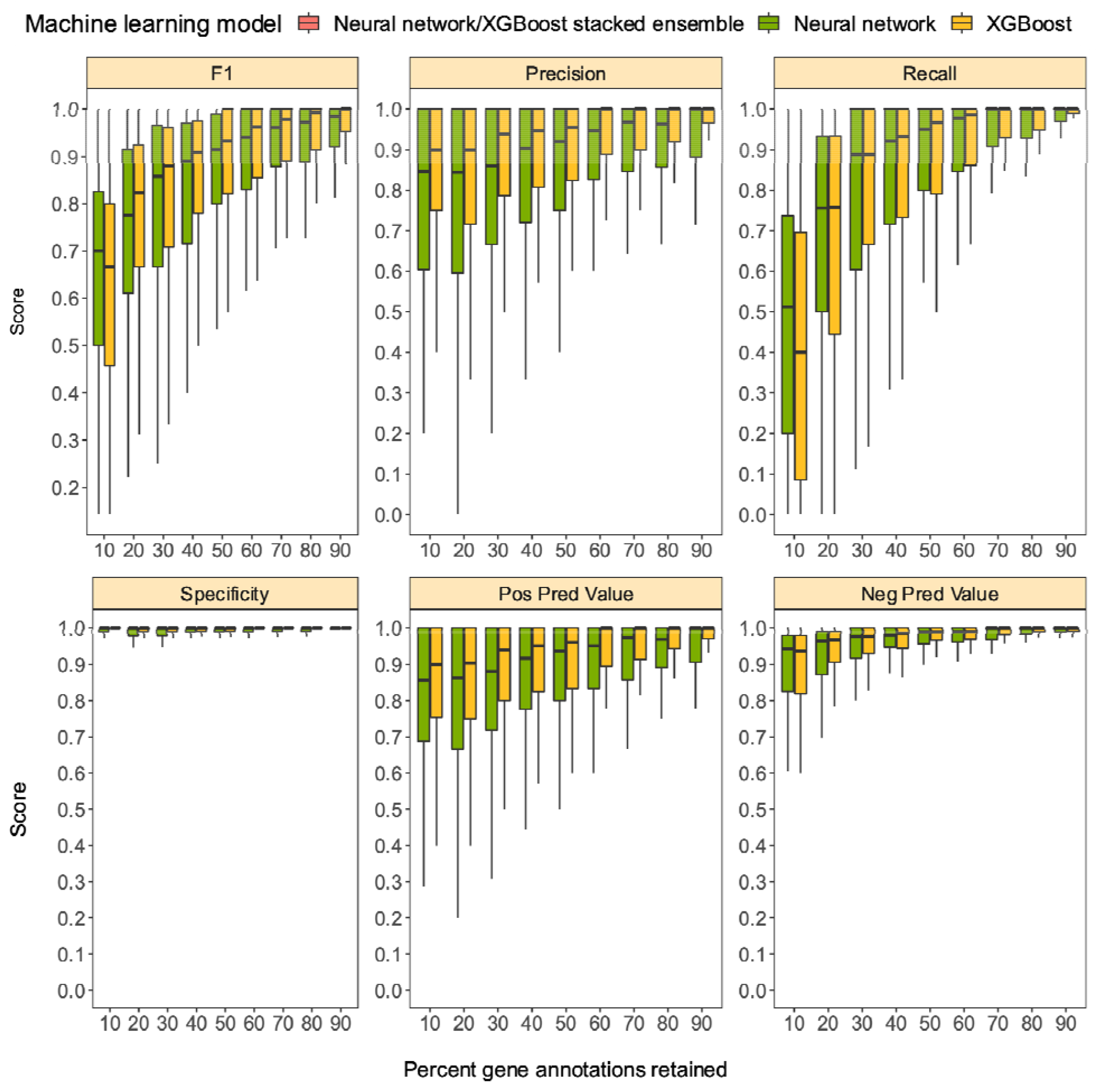
Boxplots of performance metrics of 183 neural network models on high-quality bacterial GEM MAGs (*n* = 100). Model performance metrics are for predictions on downsampled versions of GEM genome gene annotations in decreasing increments of 10% (retaining 10-90% of the annotations in each test set).

